# A universal differential expression prediction tool for single-cell and spatial genomics data

**DOI:** 10.1101/2022.11.13.516355

**Authors:** Alexis Vandenbon, Diego Diez

**Affiliations:** Institute for Life and Medical Sciences, Kyoto University, 53 Shougoin Kawahara-cho, Sakyo-ku, Kyoto 606-8507, Japan; Institute for Liberal Arts and Sciences, Kyoto University, Yoshidanihonmatsu-cho, Sakyo-ku, Kyoto 606-8501, Japan; Immunology Frontier Research Center, Osaka University, 3-1, Yamada-oka, Suita, Osaka 565-0871, Japan

## Abstract

With the growing complexity of single-cell and spatial genomics data, there is an increasing importance of unbiased and efficient exploratory data analysis tools. One common exploratory data analysis step is the prediction of genes with different levels of activity in a subset of cells or locations inside a tissue. We previously developed singleCellHaystack, a method for predicting differentially expressed genes from single-cell transcriptome data, without relying on clustering of cells. Here we present an update to singleCellHaystack, which is now a universally applicable method for predicting differentially active features: 1) singleCellHaystack now accepts continuous features that can be RNA or protein expression, chromatin accessibility or module scores from single-cell, spatial and even bulk genomics data, and 2) it can handle 1D trajectories, 2-3D spatial coordinates, as well as higher-dimensional latent spaces as input coordinates. Performance has been drastically improved, with up to ten times reduction in computational time and scalability to millions of cells, making singleCellHaystack a suitable tool for exploratory analysis of atlas level datasets. singleCellHaystack is available as an R package and Python module

## INTRODUCTION

Recent advances in single-cell and spatial omics technologies allow researchers to obtain abundance measures of transcripts and proteins, or the accessibility of genomic regions at single-cell resolution. These technologies present an unprecedented view of the heterogeneity in cell populations and their spatial distributions within tissues. However, they are also accompanied by new challenges in data analysis.

A fundamental step in exploring single-cell transcriptomics data is predicting genes that have different levels of expression in one subset of cells compared to others. Such genes are often referred to as differentially expressed genes (DEGs). Similarly, in spatial transcriptomics data, spatial DEGs are genes with altered expression in part of a tissue. In single-cell ATAC (scATAC-seq) data, differentially accessible genomic region are regions which have a higher accessibility in one group of cells compared to others. In this paper, we will use the term DEG to refer to any feature or set of features with differential levels of activity within an input space, be it the 2D or 3D space within a tissue or latent spaces – such as principal components, tSNE or UMAP – of any dimension.

The majority of single-cell DEG prediction approaches are based on two steps: 1) clustering of cells by similarity, and 2) applying statistical tests between clusters to identify DEGs ^1–7^. However, benchmark studies have reported that DEG prediction approaches for bulk RNA-seq do not perform worse than methods designed specifically for single-cell RNA-seq (scRNA-seq), and that the agreement between single-cell DEG prediction approaches is low ^8, 9^. Because the number of clusters tends to be large, a common approach is to compare the cells in each cluster against all other cells, restricting DEGs that can be detected to genes with high (or low) expression in a single cluster. For the prediction of spatial DEGs, methods have been developed that directly employ the spatial coordinates of cells (or spots or pucks) to detect genes that have non-random distributions of expression in the 2D (or 3D) space of the tissue ^10–17^. However, most existing methods do not scale well with large datasets, suffer from prohibitively long runtimes, and are limited in the spatial patterns that they can detect in practice. The development of more flexible approaches for discovering complex differential expression patterns is one of the grand challenges of this field ^18^.

We recently developed singleCellHaystack, a method that predicts DEGs based on the distribution of cells in which they are active within an input space ^19^. Our method does not rely on comparisons between clusters of cells and is applicable to both scRNA-seq and spatial transcriptomics data. Although singleCellHaystack was superior than other methods in our extensive comparison ^19^, a limitation of the implementation was that it used a hard threshold for defining genes as being either detected or not detected in each cell. Treating detection in a binary way ignores the magnitude of gene expression differences, and some differential expression patterns might be missed. Furthermore, singleCellHaystack was not able to handle sparse matrices, limiting its applicability to the ever-increasing dataset sizes.

Here, we present a new approach which addresses the above limitations. First, our new method uses continuous activity levels for predicting DEGs. Second, it uses cross-validation for choosing a suitable flexibility of splines during its modeling steps. Third, the computational time has been drastically reduced by incorporating several engineering improvements to the base code, including the use of sparse matrices. Finally, a Python implementation has been developed which enables the efficient application of singleCellHaystack to atlas level datasets with millions of cells. These improvements, together with the fact that it does not make strong assumptions about the statistical distribution of the input data, make singleCellHaystack applicable to a wide range of data types. In this manuscript we describe applications to single-cell transcriptomics, spatial transcriptomics (Visium, Slide-seqV2, HDST, and MERFISH), scATAC-seq, CITE-seq, and a large collection of bulk RNA-seq samples ^20–24^. Moreover, our approach can also be used on sets of genes (e.g., genes sharing a common annotation) for predicting differential activities of biological pathways in single-cell and spatial transcriptomics data, and to identify DEG along trajectories. Together, these results illustrate the usefulness of singleCellHaystack for exploring complex biological datasets. The new singleCellHaystack method is implemented as an R package and is available from CRAN (version 1.0.0 and higher) and GitHub, and as a Python package available from PyPI and GitHub.

## RESULTS

### Introduction to the singleCellHaystack method

Figure 1A shows a summary of the singleCellHaystack approach (version 1.0.0; see Methods for a more detailed description). In brief, singleCellHaystack requires two types of input data: 1) the coordinates of the samples (e.g., cells, spots, pucks, etc.) inside a space, which could be a 1D trajectory (e.g., pseudotime), 2D or 3D spatial coordinates, or a latent space such as principal components and 2) a matrix of numerical values reflecting the activities of features in each sample. These typically are estimates of the concentrations or RNAs or proteins, but can also be scores reflecting the accessibility of genomic regions or the average expression of sets of genes that share common functional annotations. In a first step, singleCellHaystack estimates *Q*, the distribution of samples inside the input space. It does so by measuring the local density of samples around a set of grid points. Next, for each feature *f* it estimates *P*_*f*_, the distribution of the activity of *f* inside the space employing the same grid points. This is done by weighting the density of samples around each grid point by the activity of feature *f*. The difference between each *P*_*f*_ and the reference distribution *Q* is measured by using the Kullback-Leibler divergence *D*_*KL*_^19^. The statistical significance of each *D*_*KL*_ value is estimated using randomization of the input data, and by modeling the expected distribution of *D*_*KL*_ values using splines. A suitable flexibility of the splines is determined using cross-validation. Finally, a p-value is estimated for each feature, and features with low p-values are regarded as DEGs. Because singleCellHaystack does not make comparisons between clusters, it allows detecting more complex patterns of differential activity.

**Figure 1:**
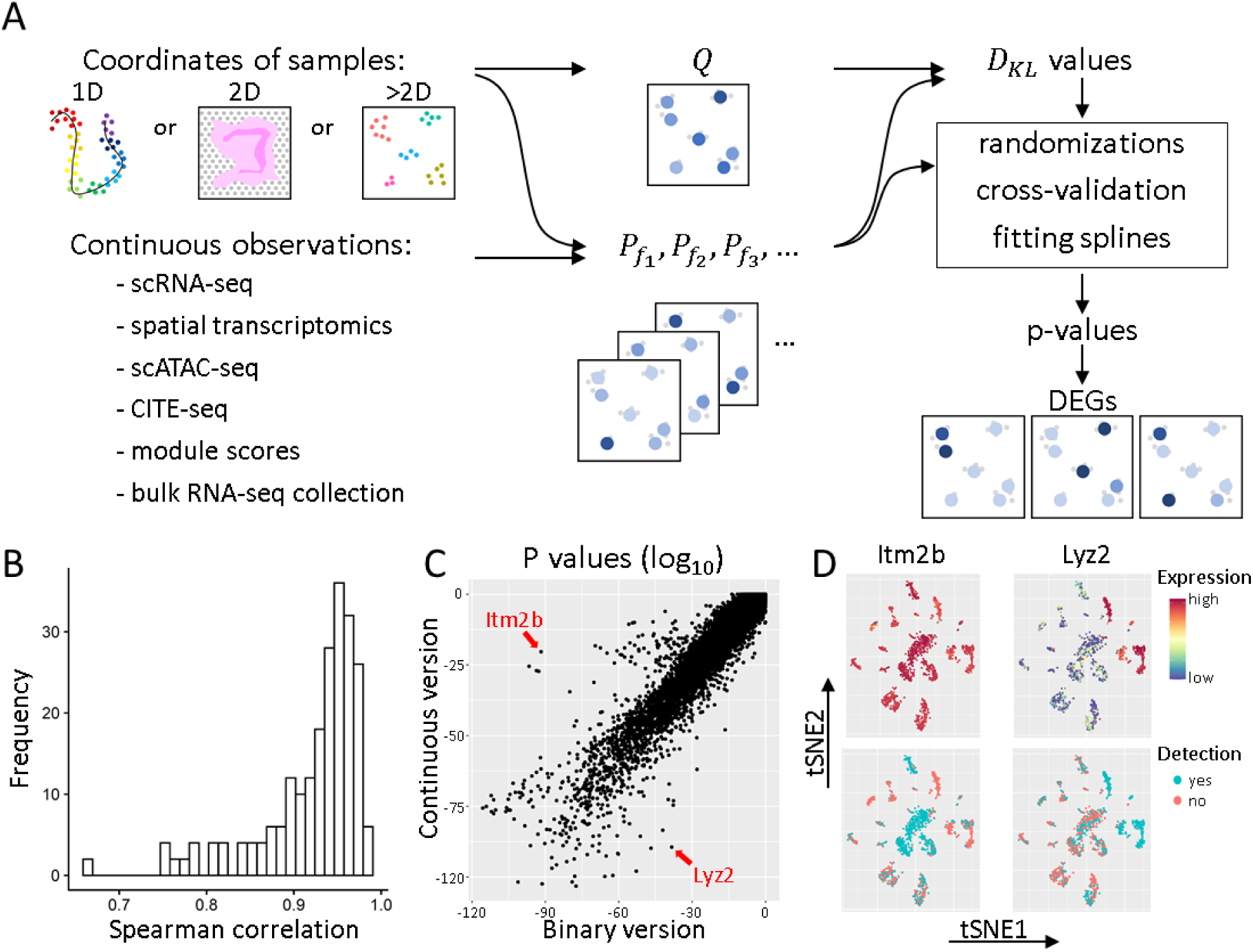
Concept of the new approach of singleCellHaystack and comparison with the previous binary approach. **(A)** A schematic overview of the new singleCellHaystack method. **(B)** Histogram of the Spearman correlation values between p-values estimated by the binary and the continuous versions of singleCellHaystack on 119 scRNA-seq datasets. **(C)** Scatter plot of p values (log_10_) estimated using the binary (X axis) and continuous (Y axis) versions of singleCellHaystack on the Tabula Muris Lung dataset. Two genes with large discrepancies in p-values are indicated. **(D)** tSNE plots showing the genes indicated in **(C)**, expression values as used by the continuous version (top) and detection levels as used by the binary version of singleCellHaystack (bottom).

Our previous method (available in singleCellHaystack version 0.3.2) treated the expression of genes as binary (i.e. either expressed or not) using a hard threshold ^19^. This new method (found in singleCellHaystack version 1.0.0 and higher) uses continuous values, reflecting the magnitude of the activity of each feature around each grid points, allowing it to identify more nuanced patterns of expression, and extending the application to features other than gene expression levels. To investigate differences in DEGs returned by the new continuous version and the original binary version of singleCellHaystack, we applied both versions on 119 scRNA-seq datasets of Tabula Muris and Mouse Cell Atlas ^25, 26^. Although both methods returned generally consistent results (the average Spearman correlation between log p-values of the binary and the continuous version was 0.92; Fig. 1B), in each dataset we also observed large discrepancies caused by the usage of the hard threshold in the binary version. For example, both versions returned highly consistent results on the Tabula Muris lung tissue dataset (Spearman correlation 0.95; Fig. 1C), but several genes were judged to have differential expression by one version and not by the other. Two examples are *Itm2b* and *Lyz2* (Fig. 1C-D). *Itm2b* has stable expression levels across the cell clusters in the dataset. In contrast, *Lyz2* has high expression in a few subsets of cells, with lower expression in most others. In the binary version, the relatively small differences in expression of *Itm2b* become exaggerated because of the use of a hard threshold. On the other hand, for *Lyz2* the binary version dilutes the differential expression pattern because some cells with low expression also exceed the hard threshold. This leads to *Itm2b* being regarded as a top-scoring DEG by the binary version but not by the continuous version (ranked 42^nd^ by the binary version; 2872^nd^ by the continuous version), and the opposite result for *Lyz2* (ranked 1014^th^ by the binary version; 42^nd^ by the continuous version). Other examples are shown in Supplementary Fig. S1.

In addition to the changes to the *D*_*KL*_ computation, we made improvements to other parts of the implementation, including efficient use of sparse matrices that resulted in shorter runtimes (Supplementary Fig. S2). The continuous version was faster than the binary version on all 119 scRNA-seq Tabula Muris and Mouse Cell Atlas datasets, with an average 37.8% reduction in runtime. In addition, we implemented singleCellHaystack in Python (https://github.com/ddiez/singleCellHaystack-py) enabling broader usability and the application to very large datasets. To show this we applied the Python version to a scRNA-seq dataset with 4 million cells from human fetal tissues ^27^ (Supplementary Figure 3). This analysis took around 165 minutes to finish on a workstation with 28 cores and 768 GB of physical memory indicating that singleCellHaystack scales to atlas-level datasets with millions of cells.

### Application to spatial transcriptomics data and comparison with existing methods

We applied several spatial DEG prediction methods on spatial transcriptomics data of several platforms (MERFISH, 10x Visium, Slide-seqV2, and HDST; see Table 1). The methods compared were singleCellHaystack, SPARK, SPARK-X, Seurat’s FindSpatiallyVariableFeatures using the Moran’s I and mark variogram approaches, MERINGUE, and Giotto’s binSpect using the kmeans and the rank approaches ^12, 15–17, 19, 28, 29^. We first ran all methods on the top 1,000 highly variable genes (HVGs) in each dataset and recorded their runtimes. Unfortunately, most methods do not scale well with increasing dataset size (Fig. 2A), or failed to run on the larger datasets. SPARK-X was in general the fastest method, followed by singleCellHaystack. Runtimes of singleCellHaystack are not solely a function of the number of spots (or cells, pucks) in the data, but also of the sparsity of the data. Because of this, runs on HDST datasets (which have a lot more zeroes) took less time than runs on Slide-seqV2 datasets of similar sizes.

**Figure 2:**
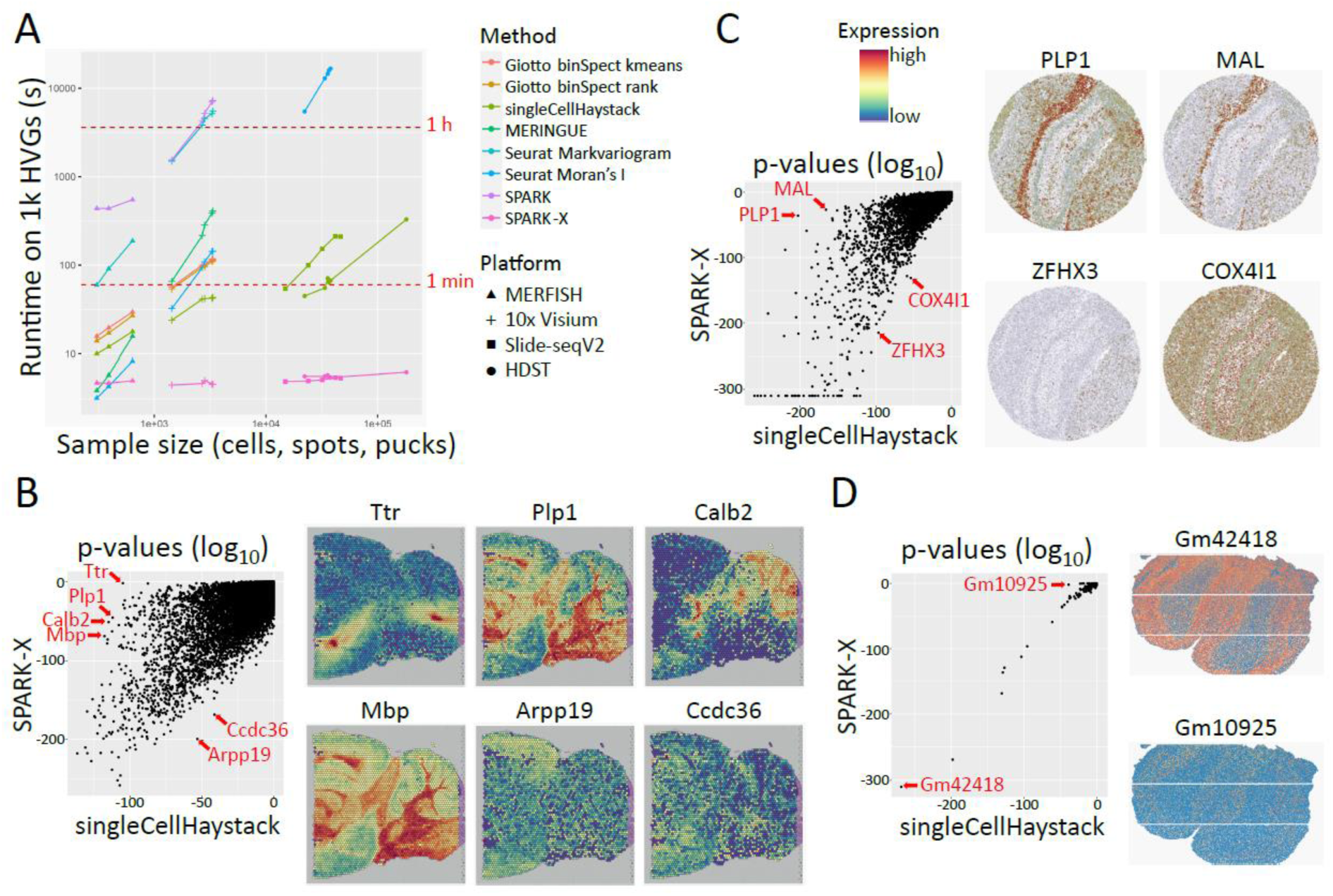
Applications to spatial transcriptomics data. **(A)** Comparison of runtime of several spatial DEG prediction methods applied on 1,000 HVGs of datasets of different platforms. For fair comparison, SPARK and SPARK-X were run on 1 core. Applications which failed to return results are not shown. **(B-D)** Example comparisons of the results of singleCellHaystack and SPARK-X. For each comparison, a scatterplot of p values (log_10_) is shown on the left, and examples of DEGs are shown on the right. Shown examples are for mouse posterior brain (10x Visium, dataset “posterior1”) **(B)**, mouse hippocampus (Slide-seqV2) **(C)**, and mouse olfactory bulb (HDST) **(D)**.

**Table 1:** Overview of spatial transcriptomics datasets.

| ID | Name | Platform | Species | Tissue / origin | Reference |
| --- | --- | --- | --- | --- | --- |
| 1 | Anterior1 | Visium | mouse | brain | 10x Genomics |
| 2 | Anterior2 | Visium | mouse | brain | 10x Genomics |
| 3 | Posterior1 | Visium | mouse | brain | 10x Genomics |
| 4 | Posterior2 | Visium | mouse | brain | 10x Genomics |
| 5 | Kidney | Visium | mouse | kidney | 10x Genomics |
| 6 | Stickels_hippo1 | Slide-seqV2 | mouse | hippocampus | 20 |
| 7 | Stickels_hippo2 | Slide-seqV2 | mouse | hippocampus | 20 |
| 8 | Stickels_embryo | Slide-seqV2 | mouse | embryo | 20 |
| 9 | Stickels_olfactory | Slide-seqV2 | mouse | olfactory bulb | 20 |
| 10 | Stickels_cortex | Slide-seqV2 | mouse | cortex | 20 |
| 11 | Vickovic_CN13_D2 | HDST | mouse | olfactory bulb | 21 |
| 15 | Vickovic_CN24_D1 | HDST | mouse | olfactory bulb | 21 |
| 16 | Vickovic_CN24_E1 | HDST | mouse | olfactory bulb | 21 |
| 12 | Vickovic_CN21_C1 | HDST | human | breast cancer | 21 |
| 13 | Vickovic_CN21_D1 | HDST | human | breast cancer | 21 |
| 14 | Vickovic_CN21_E2 | HDST | human | breast cancer | 21 |
| 17 | Xia_B1 | MERFISH | human | osteosarcoma | 22 |
| 18 | Xia_B2 | MERFISH | human | osteosarcoma | 22 |
| 19 | Xia_B3 | MERFISH | human | osteosarcoma | 22 |

For a genome-wide comparison on all genes of each datasets, we restricted ourselves to the two fastest methods (singleCellHaystack and SPARK-X). We applied both methods on each dataset and plotted the returned p-values (log_10_ values) (Fig. 2B-D, Supplementary Fig. S4-S7). In several datasets, we observed that singleCellHaystack was able to pick up clear DEGs which were missed by SPARK-X. For example, in the mouse posterior brain 10x Visium dataset, top-scoring DEGs by singleCellHaystack included *Mbp*, *Calb2*, *Plp1*, and *Ttr* (Fig. 2B), all of which show clear spatially differential expression patters, yet were not among top-scoring DEGs predicted by SPARK-X. A striking example is the highly concentrated expression of *Ttr* (transthyretin) in 2 locations in the posterior brain. singleCellHaystack regards *Ttr* as a top-scoring DEG (p value 1.1 x 10^-105^; ranked 58^th^ out of 16,596 genes), but SPARK-X does not (p-value 0.0087; ranked 11,881^st^). In contrast, genes that are top-scoring according to SPARK-X but not singleCellHaystack are rare. Two such genes are Arpp19 (ranked 18^th^ by SPARK-X; 983^rd^ by singleCellHaystack) and Ccdc36 (ranked 73^rd^ by SPARK-X; 1,585^th^ by singleCellHaystack) in the same posterior brain dataset. Although both genes exhibit some spatial expression pattern, it is relatively weak compared to *Ttr* and other genes which are missed by SPARK-X. Similar results were seen in other datasets from posterior brain, anterior brain and kidney (Supplementary Figure S4).

In Slide-seqV2 datasets, too, top-scoring DEGs picked up by singleCellHaystack but missed by SPARK-X were relatively common. For example, in the mouse hippocampus sample, several genes were top-ranking DEGs according to singleCellHaystack but not SPARK-X, including *Plp1* (ranked 20^th^ by singleCellHaystack vs 1,139^th^ by SPARK-X) and *Mal* (ranked 55^th^ by singleCellHaystack vs 1,543^rd^ by SPARK-X), all showing strong spatial expression patterns (Fig. 2C, Supplementary Figures S5). In contrast, there were no genes that were top-scoring according to SPARK-X but missed by singleCellHaystack. Two examples of genes that were relatively higher-scoring for SPARK-X than for singleCellHaystack are *Zfhx3* (378^th^ by singleCellHaystack vs 64^th^ by SPARK-X) and *Cox4i1* (1123^rd^ by singleCellHaystack vs 183^rd^ by SPARK-X). Although both *Zfhx3* and *Cox4i1* show some degree of spatially differential expression, the tendency is weaker than those of *Plp1* and *Mal*.

In HDST datasets, singleCellHaystack and SPARK-X returned highly consistent results (Fig. 2D and Supplementary Fig. S6). For example, gene Gm42418 was the top-scoring DEG according to both methods in one mouse brain sample (Fig. 2D). Only 1 gene was found to have a discrepant result: Gm10925 was ranked 13^th^ by singleCellHaystack (p value 2.7 x 10^-39^) but only 105^th^ by SPARK-X (p value 0.013).

Finally, we compared both methods using three MERFISH datasets (Supplementary Figures S7) ^22^. Although the differential patterns of expression are visually less clear, singleCellHaystack was able to pick up genes for which cells with high/low expression are located proximally in space.

In summary, top-scoring DEGs predicted by singleCellHaystack include clear spatial DEGs, including cases that are missed by SPARK-X (Fig. 2B-D). The authors of SPARK-X noted that the assumptions made by SPARK-X are likely to be not optimal in detecting certain expression patterns ^16^. In our results, SPARK-X appears to work well on gradually changing patterns of expression, but suffers on patterns with abrupt differences between neighboring locations, exemplified by *Ttr* in Fig. 2B or *PLP1* in Fig. 2C.

So far, we have described applications to the multidimensional PC space of scRNA-seq data and 2D spatial coordinates of various spatial transcriptomics technologies. However, singleCellHaystack makes few assumptions about the underlying data distributions (distribution of read counts or UMIs, etc.). Because of the versatility of the density distribution and relative entropy approach on which singleCellHaystack is based, it is applicable to many other data types and input spaces. In the next sections we illustrate this general applicability using examples on scRNA-seq trajectory (1D) data, CITE-seq data, scATAC-seq data, a large collection of bulk RNA-seq data, and on gene set activity data.

### Predicting DEGs along trajectories

We applied singleCellHaystack to 1D projections from trajectory pseudotime inference. To this end we used the thymus dataset from the Tabula Muris ^25^, which contains data from developing thymocytes, progressing from a double negative (Cd4^-^Cd8^-^) through a double positive (Cd4^+^Cd8a^+^) stage into mature naive T cells characterized as single positive (i.e., either Cd4^+^ or Cd8a^+^). We processed the 10x Genomics Chromium data using the standard pipeline with Seurat ^30^ and then used monocle3 ^31^ to order cells from the double negative cluster to the single positive clusters (Fig. 3A). We used this pseudotime ordering (a 1D space) as input with singleCellHaystack to identify DEGs with biased expression along this trajectory. To characterize the patterns and dynamics associated with the changes in expression we clustered the DEGs into 6 modules. Figure 3B shows the mean expression of the top-scoring genes in each module along the trajectory, whereas Figure 3C shows the top 10 genes per module. These results indicate that singleCellHaystack is able to identify patterns of gene expression changes along trajectories.

**Figure 3:**
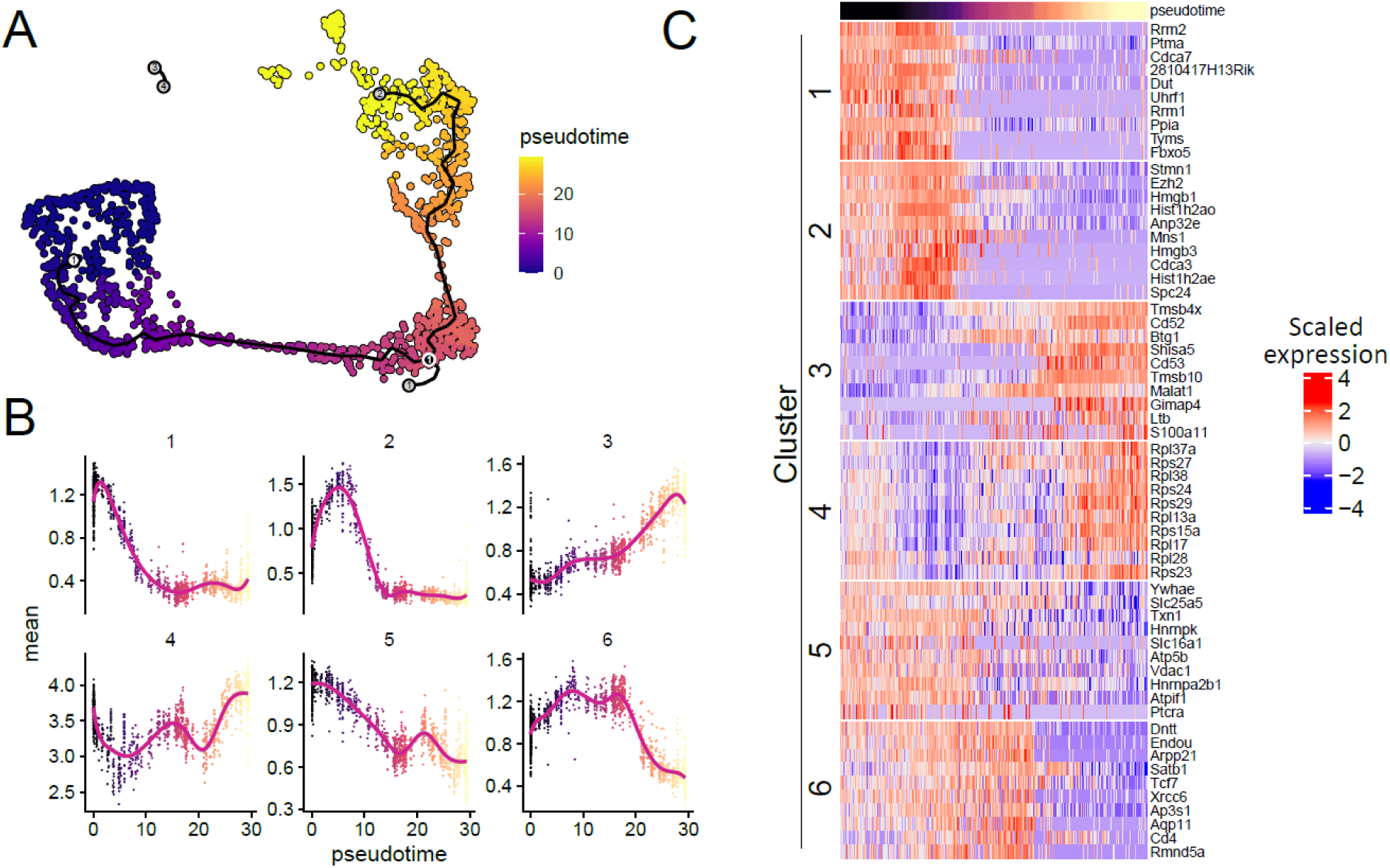
Application of singleCellHaystack to the prediction of DEGs along a trajectory. **(A)** UMAP plot of a Tabula Muris thymus dataset indicating the monocle3 trajectories and pseudotime. **(B)** Top scoring DEGs predicted by singleCellHaystack were clustered into modules. For each module the average expression of all genes at each pseudotime value is shown, indicating different patterns of expression changes along the trajectory. **(C)** For each module in panel **(B)**, the expression of the top 10 genes along the trajectory is shown in a heatmap.

### Applications to CITE-seq, scATAC-seq, and bulk RNA-seq data

Our method is not restricted to single-cell transcriptome data, but can be used with any numerical data. Here we demonstrate this by applying singleCellHaystack to CITE-seq, scATAC-seq and bulk RNA-seq data.

To show singleCellHaystack applications to single-cell protein measurements we used a human peripheral blood mononuclear cell (PBMC) dataset containing the whole transcriptome, and the expression of more than 200 proteins ^30^. We calculated PCA and UMAP coordinates, and cell clusters based on the expression of proteins and ran singleCellHaystack using 50 PCs and the protein counts. Figure 4A shows the UMAP plot with the clusters, together with the top 8 proteins identified by singleCellHaystack.

**Figure 4:**
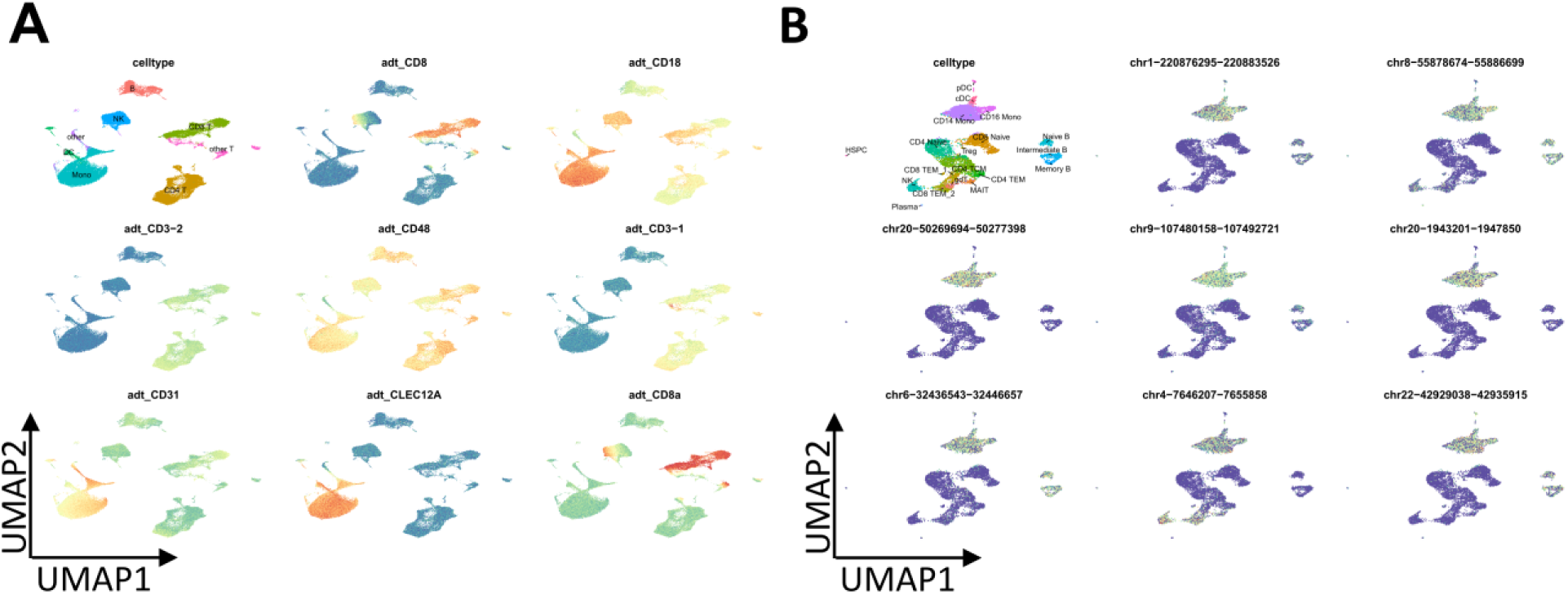
Example application to CITE-seq and scATAC-seq data. **(A)** A UMAP plot of the CITE-seq data with cell type annotations is shown (top left) together with the top 8 high-scoring genes predicted by singleCellHaystack. **(B)** A UMAP plot of the scATAC-seq data with cell types annotations is shown (top left) together with the top 8 high-scoring differentially accessible genomic regions predicted by singleCellHaystack.

We also applied singleCellHaystack to a single-cell multiome (i.e., RNA and ATAC) dataset from human PBMCs downloaded from the 10x Genomics website (see Methods). We use the Signac package to process the RNA and ATAC counts. For ATAC we calculated a Latent Semantic Index (LSI) embedding and used it, with the peak counts, to identify differential accessibility regions. Figure 4B shows the UMAP plot derived from the LSI, together with the top 8 regions identified by singleCellHaystack.

Another possible application is on large numbers of bulk RNA-seq samples. Here, as an example, we applied singleCellHasytack on a collection of 1,958 RNA-seq samples obtained from various parts of the mouse brain. singleCellHaystack successfully predicted DEGs with differential expression in subsets of the samples (Supplementary Fig. S8) ^23^.

### Predicting differentially active gene sets

Because singleCellHaystack makes few assumptions about the input data, it is not limited to applications to UMI or read count data, but can be used with any quantitative data associated with the samples. As an illustration, we applied singleCellHaystack to module scores as computed by Seurat, which reflect the general activity of a set of genes. Here, as sets of genes, we used genes associated with 292 pathways as defined in BioCarta by msigdbr ^32^. For a number of spatial transcriptomics datasets, we calculated the module scores of each gene set in each Visium spot, and used singleCellHaystack to predict gene sets with highly non-random spatial distributions. Examples of high-scoring gene sets in mouse anterior and posterior brain and kidney tissue are show in Figure 5. For each dataset a variety of patterns was found, reflecting how different pathways are active in different parts of the tissues.

**Figure 5:**
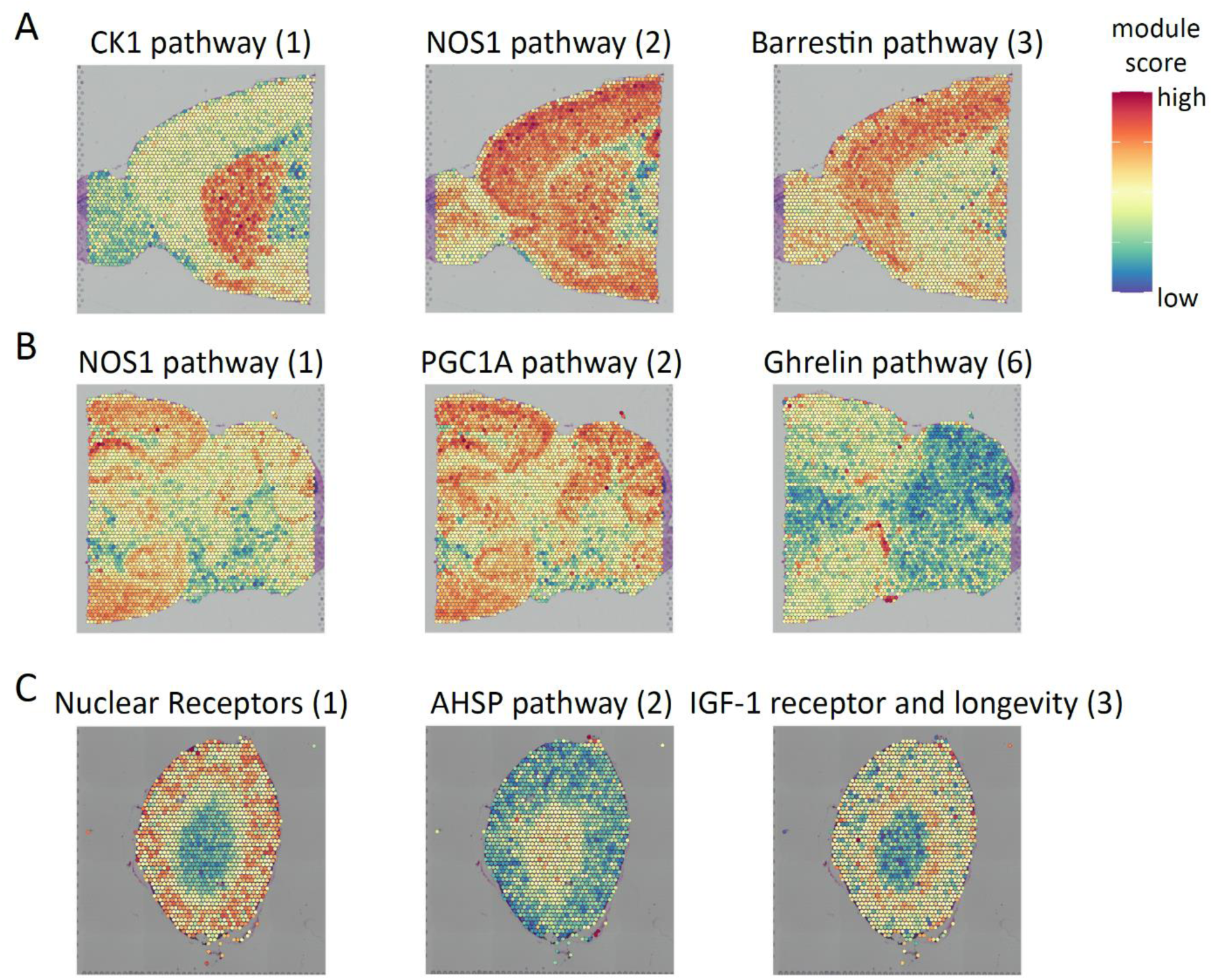
Application of singleCellHaystack to sets of genes. We applied singleCellHaystack on the module scores of sets of genes associated with 292 BioCarta pathways. Examples of high-scoring BioCarta pathways are shown in three spatial transcriptomics datasets. Numbers in parentheses represent the rank of the p-value of the pathway (e.g. 1 indicates the most significant pathway). **(A)** in mouse anterior brain, **(B)** in mouse posterior brain, and **(C)** in mouse kidney.

## DISCUSSION

In this manuscript we have presented a universally applicable tool for differential expression prediction in single-cell and spatial genomics data. This version of singleCellHaystack is an important improvement compared to existing DEG prediction tools and compared to the original implementation of singleCellHaystack. The original implementation required binary detection data as input (i.e. genes were treated as either detected or not in each cell), thus ignoring more subtle differences in expression between subsets of cells. In this new version this issue has been addressed, and singleCellHaystack now uses continuous activity levels of features to detect DEGs. This enables singleCellHaystack to be used with any kind of continuous measurements, whether RNA or protein levels, chromatin accessibility or gene ontology scores. Furthermore, we showed that we can use singleCellHaystack with any kind of cell coordinates, whether they are physical spatial locations, PCA embeddings or pseudotime ranking.

Single-cell genomic datasets are rapidly increasing in numbers and in size, making it more challenging to perform exploratory analyses, including the identification of DEGs. Our new implementation of singleCellHaystack is significantly more efficient and faster than the original version, making it possible to analyzed large datasets in a few minutes. For example, the Mouse Organogenesis Cell Atlas dataset with over 100 thousand cells took ∼45 minutes to finish with the original version, whereas it takes around 5 minutes to finish with the new one (not shown). Furthermore, our new Python implementation enables efficient identification of DEGs for atlas level datasets with millions of cells.

For spatial transcriptomics the fastest method available is SPARK-X. The short runtimes are accomplished by, among others, making several assumptions about the input data ^16^. Possibly because of these assumptions, SPARK-X failed to identify several clear DEGs in spatial transcriptomics datasets, when compared to singleCellHaystack (see for example Fig. 2B). In contrast, we found no clear DEGs that were predicted by SPARK-X but not by singleCellHaystack. The better sensitivity of singleCellHaystack makes it the best alternative, despite relatively longer runtimes.

Different methods for the identification of DEGs are being used depending on the technology. For example, Wilcoxon rank sum tests and t-tests are used to identify DEG between groups of cells. Moran’s I and other methods are used to identify DEGs in spatial transcriptomics and trajectory analyses. In this paper we show that singleCellHaystack is not restricted by the technology or by the particularities of features that were measured, nor by the type of coordinates that were used as input. This makes singleCellHaystack a universal tool for the identification of DEGs.

Despite its advantages, singleCellHaystack has a few weak points. One is that comparisons between multiple conditions (e.g., wild-type and knockout) cannot easily be conducted. We hope to expand singleCellHaystack to include methods for such comparisons in the future. Secondly, for better or worse, current scRNA-seq data analyses are often cluster-oriented. The clustering of cells is a convenient tool for summarizing complex datasets and for performing additional downstream tests. Compared to cluster-based DEG prediction approaches, it is not as straightforward to incorporate the results of singleCellHaystack into a cluster-oriented scRNA-seq analysis. However, other fields of genomics, such as spatial transcriptomics, are less focused on clustering, and not all scRNA-seq datasets can be easily summarized by clustering. We believe that a method like singleCellHaystack, which can detect complex patterns without being restricted by clusters, will play a valuable role in future exploratory analysis.

## MATERIALS AND METHODS

### singleCellHaystack methodology

For a detailed description of the original singleCellHaystack implementation (version 0.3.2) we refer to Vandenbon and Diez ^19^. In brief, singleCellHaystack uses the distribution of cells inside an input space to predict DEGs. First, it infers a reference distribution *Q* of all cells in the space by estimating the local density of cells surrounding a set of grid points in the space. In a next step, the original singleCellHaystack estimated the distribution of the cells in which a gene *G* is detected (distribution *P*(*G* = *T*)) and not detected (distribution *P*(*G* = *F*)). The Kullback-Leibler divergence (*D*_*KL*_) was used to compare *P*(*G* = *T*) and *P*(*G* = *F*) to the reference distribution *Q*. The statistical significance of each gene’s DKL was evaluated using random sampling.

The updated version of singleCellHaystack (version 1.0.0) includes several improvements. The main improvement is that singleCellHaystack no longer treats expression in a binary way (i.e. detected or not detected), but uses continuous values (see Steps 3-4 below). Secondly, we updated the modeling of *D*_*KL*_ values using splines. In the new implementation, we use cross-validation to select a suitable flexibility of the splines (see Step 5). The new implementation also accepts input data as sparse matrices, and a Python implementation has been made available. Below follows a more detailed description of the singleCellHaystack version 1.0.0 methodology.

### Step 1: Setting parameters

The main inputs to singleCellHaystack are the coordinates of samples inside an input space, and the observations in each sample. Here, samples include single cells, spots, pucks or even bulk samples, depending on the platform used. The input space could be the 2D or 3D space in a tissue or a latent space after typical dimensionality reduction (e.g. first principal components of a scRNA-seq dataset), or the 1D coordinates of samples along a trajectory (pseudotime). Coordinates of samples in *d*-dimensional space will be denoted as *s* ∈ *R*^*d*^. By default, the coordinates in each dimension are rescaled to mean 0 and standard deviation 1. Observations might be gene expression, chromatin accessibility, gene set module scores, etc. and will be denoted as *y*.

Steps 2-4 involve estimating the distribution of samples inside the *d*-dimensional input space by estimating the local density of samples around a set of grid points using a Gaussian kernel. By default, the coordinates of grid points are decided by running k-means clustering on the sample coordinates *s* and then using the resulting centroids as grid points. Note that the goal of this step is not to obtain clusters of cells, but merely to obtain suitable grid points. This approach tends to result in grid points being roughly uniformly spread over the subspace of the input space where samples are located. By default, singleCellHaystack uses *g*=100 grid points (option grid.points). This number can be reduced (e.g. when the number of samples is low) or increased (for highly heterogeneous datasets or when the number of samples is high) as needed. We have shown before that results are stable w.r.t. the number of grid points ^19^. Alternatively, the user can specify the coordinates of the grid points to use (option grid.coord), or use seeding, as used in k-means++ clustering, as described before (grid.method="seeding") ^19^.

The bandwidth ℎ of the Gaussian kernel is set as before ^19^. For each sample, the Euclidean distance to the closest grid point is calculated, and ℎ is defined as the median of those distances. Normalized distances between samples and grid points are subsequently defined as the Euclidean distances divided by the bandwidth ℎ. The density contribution *d*_*i*,*j*_ of each sample *i* to each grid point *j* is calculated as:

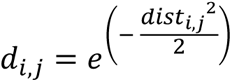

where *dist*_*i*,*j*_ is the normalized distance between sample *i* and grid point *j*.

### Step 2: Estimating reference distribution Q

The reference distribution *Q* = (*Q*_1_, …, *Q*_*g*_) of all *n* samples in a dataset is estimated as described before ^19^. In brief, the density of cells around grid point *j* is calculated as the sum of all *d*_*i*,*j*_ values:

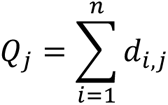

After this, *Q* is normalized to sum to unity.

### Step 3: Estimating *P*_*f*_ distributions

Whereas the original version of singleCellHaystack treated observations in a binary way (a gene is either detected or not detected in each cell), this updated version of singleCellHaystack addresses this weak point and treats observations in a continuous manner. To do so, the distribution *P*_*f*_ = (*P*_*f*,1_, …, *P*_*f*,*g*_) of feature *f* in the input space is calculated as follows:

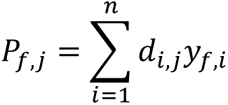

where *d*_*i*,*j*_ is the density contribution of sample *i* to grid point *j*, and *y*_*f*,*i*_ is the activity of feature *f* in sample *i*. *P*_*f*,*j*_ is therefore the sum of density contributions of samples to grid point *j* weighted by the activity of *f*. Subsequently, *P*_*f*_ is normalized to sum to unity.

### Step 4: Estimating the Kullback-Leibler divergence of feature *f*, *D*_*KL*_(*f*)

The divergence of feature *f*, *D*_*KL*_(*f*), is calculated as follows:

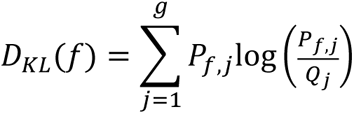

This approach is simpler than the original version, because no distinction needs to be made between samples in which a feature was detected or not detected ^19^. If the activity of feature *f* does not show a biased distribution, and approximately follows the reference distribution *Q*, then *D*_*KL*_(*f*) is close to 0. As the discrepancy with the reference distribution *Q* increases, the value of *D*_*KL*_(*f*) also increases.

### Step 5: Estimating the significance of *D*_*KL*_(*f*)

In a final step, singleCellHaystack evaluates the statistical significance of *D*_*KL*_(*f*) values by comparing them to randomized data. In principle, it would be possible to generate many randomly shuffled sets of activity values of each feature *f*, and use these to estimate a null distribution of randomized *D*_*KL*,*random*_(*f*) values. However, doing this for each feature would be prohibitively time-consuming. Instead, singleCellHaystack uses the following approach.

First, singleCellHaystack calculates the coefficient of variation (CV = standard deviation / mean) of each feature *f*. Features are ordered by CV, and a subset of features (100 by default) that are spread evenly over the range of CV values is selected. These features are used for making randomly permutated datasets (100 by default for each selected feature) based on which *D*_*KL*,*random*_(*f*) values are calculated.

For each randomized feature *f*, the *log*(*D*_*KL*,*random*_(*f*)) values follow an approximately normal distribution. This allows us to use their mean and standard deviation to estimate p-values of actually observed *D*_*KL*_(*f*) values. Moreover, we can use CV values as predictor of the mean and standard deviations of *log*(*D*_*KL*,*random*_(*f*)) values. In singleCellHaystack, by default we model the mean and standard deviation of the *log*(*D*_*KL*,*random*_(*f*)) values in function of *log*(CV) values using natural cubic splines. Splines are trained using function ns in the splines R package. A suitable degree of freedom (between 1 and 10) is decided using 10-fold cross-validation.

Alternatively, B-splines can be used, using function bs. In this case, a suitable degree (between 1 and 5) and degree of freedom (between 1 and 10) is decided in the same way.

Using the splines, the expected mean and standard deviation of *log*(*D*_*KL*,*random*_(*f*)) are predicted for every feature, in function of its CV, and based on that the corresponding p-value of *D*_*KL*_(*f*) using the pnorm function in R.

### Application to Tabula Muris and Mouse Cell Atlas scRNA-seq datasets

We obtained the Tabula Muris data from https://figshare.com/articles/dataset/Robject_files_for_tissues_processed_by_Seurat/5821263 (version 3) ^25^. For the Mouse Cell Atlas data, we downloaded file MCA_BatchRemove_dge.zip from https://figshare.com/articles/MCA_DGE_Data/5435866 ^26^. This data has been treated to reduce batch effects. Data from both Tabula Muris (32 datasets) and Mouse Cell Atlas (87 datasets) were processed using the Seurat R package (version 4.0.0) ^30^. We used the same pipeline for the processing and normalization of all datasets: genes detected in less than 3 cells and cells with fewer than 100 detected genes were removed. After this initial filtering, cells with extreme UMI counts (bottom 1 percentile and top 1 percentile), or extreme numbers of detected genes (bottom 1 percentile and top 1 percentile), or with a high fraction of mitochondrial reads (>10%) were removed. The data for the remaining cells of each dataset was normalized (NormalizeData, default settings), scaled (ScaleData, regressing out the UMI count and mitochondrial fraction), and highly variable genes (HVGs) were detected (FindVariableFeatures, default settings). The HVGs were used for principal component analysis (PCA), and the 20 first principal components (PCs) were used for further dimensionality reduction (t-SNE and UMAP) and for clustering of cells (FindNeighbors and FindClusters, using 20 PCs and otherwise default settings). Both the binary (version 0.3.2) and updated (version 1.0.0) singleCellHaystack were applied on the first 20 PCs of each dataset. For the binary version, in each dataset, for each gene, the median expression level of each gene was used as a threshold to define detection. The detection data was used as input. For the updated version, the continuous expression levels were used as input.

### Application to large scRNA-seq dataset

We downloaded scRNA-seq expression data for 4,062,980 cells and 35,686 transcripts from fetal tissues ^27^. For the analysis we used our Python implementation of singleCellHaystack (https://github.com/ddiez/singleCellHaystack-py) using PCA coordinates with 50 components.

### Application to trajectory analysis

We applied singleCellHaystack to pseudotime projection on the Tabula Muris thymus data ^25^. In this dataset the development of T cells can be followed from double negative (CD4^-^CD8^-^), through double positive (CD4^+^CD8^+^) and into mature, single positive (CD4^+^CD8^-^ and CD4^-^CD8^+^) T cells. To identified the differentiation trajectory we used monocle3 ^31^. Briefly, the data was first processed using the standard Seurat pipeline (see above), except that 30 PCs were used to calculate UMAP coordinates. We converted the Seurat object into a cell_data_set object with the SeuratWrappers package (https://github.com/satijalab/seurat-wrappers). Then monocle3 was used to calculate clusters and partitions using the UMAP coordinates with the function cluster_cells. Next, the principal graph is learned using the learn_graph function, and cell were ordered selecting as root the node in the graph starting in the cluster of double negative cells.

We used singleCellHaystack using the pseudotime coordinates. We selected the top 1,000 predicted DEGs and clustered them into modules using kmeans, using k=6.

### Application to CITE-seq

Single-cell data from human peripheral blood mononuclear cells (PBMC) data was downloaded from https://atlas.fredhutch.org/nygc/multimodal-pbmc/. This dataset contains information about the expression of 228 immune marker proteins on over 200k cells. As input to singleCellHaystack we used the protein based PCA coordinates (50 PCs), and the normalized expression levels included in the downloaded data, which was processed as described here ^30^.

### Application to scATAC-seq

Single-cell multiome (RNA + ATAC) data from human PBMC was downloaded from 10x web site (https://support.10xgenomics.com/single-cell-multiome-atac-gex/datasets/1.0.0/pbmc_granulocyte_sorted_10k). The raw data (fragments and peak information from cellranger) were processed with Signac ^33^, following the workflow described here: https://satijalab.org/seurat/articles/weighted_nearest_neighbor_analysis.html#wnn-analysis-of-10x-multiome-rna-atac-1. Briefly, the expression and chromatin accessibility peak information data were loaded into a Seurat object. For the peaks, the information about genomic ranges was obtained using the function GetGRangesFromEnsDb with the Bioconductor package EnsDb.Hsapiens.v86 (http://bioconductor.org/packages/EnsDb.Hsapiens.v86/). Cells were filtered to have less than 20% of mitochondrial counts, RNA counts between 1,000 and 25,000 and ATAC counts between 5x10^3^ and 7x10^7^. For the RNA data the SCTransform pipeline was used, and UMAP coordinates calculated using 50 PCs. For the ATAC counts, first term-frequency inverse-document-frequency was calculated with RunTFIDF. Top features were selected with FindTopFeatures and min.cutoff="q0". Then, a Latent Semantic Index (LSI) embedding was calculated with RunSVD. ATAC based UMAP was constructed from dimensions 2 to 50 from LSI. singleCellHaystack was run using LSI embedding and ATAC peak counts.

### Applications to spatial transcriptomics datasets

We obtained and processed data for the following four platforms (Table 1).

### Visium platform data

Data for mouse kidney and brain were obtained through the SeuratData R package ^34^. We filtered out mitochondrial genes and genes with non-zero counts in less than 10 spots. Data was normalized using the Seurat R package (function NormalizeData, with default parameters).

### Slide-seqV2 data

We obtained the data from the Broad Institute Single Cell Portal (accession number SCP815) ^20^. We filtered out mitochondrial genes and genes with non-zero counts in less than 10 spots, as well as pucks with less than 100 reads in total. Data was normalized using the Seurat R package (function NormalizeData, with default parameters).

### HDST data

We obtained the data from the Broad Institute Single Cell Portal (accession number SCP420) ^21^. We filtered out mitochondrial genes and genes with non-zero counts in less than 10 spots. Data was normalized using the Seurat R package (function NormalizeData, with default parameters).

### MERFISH data

RNA counts and cell position information was obtained from Xia et al. (Datasets S12 and S15) ^22^. We filtered out genes with non-zero counts in less than 10 spots, as well as cells with less than 6,000 detected genes. Data was normalized using the Seurat R package (function NormalizeData, with default parameters).

We applied the following methods on each of the spatial datasets: the updated version of singleCellHaystack, SPARK, SPARK-X, Seurat’s FindSpatiallyVariableFeatures function using Moran’s I and mark variogram approaches, MERINGUE, and Giotto’s binSpect using the kmeans and the rank approaches ^12, 15–17, 19, 28, 29^. Because several methods have long runtimes or returned errors on large datasets (Fig. 2A), we limited the analysis to the top 1,000 highly variable genes (detected using function FindVariableFeatures). In addition, we applied singleCellHaystack and SPARK-X on all genes in the datasets. To each method, we gave as input the same 2D spatial coordinates of the samples, along with the expression data (counts or processed data as needed). Each method was run with default parameter settings on the 1,000 HVGs (without other pre-filtering steps), and the number of cores used was set to 1, to make the comparison of runtimes fair. Method-specific normalization steps were not included in the runtimes. SPARK was applied using CreateSPARKObject (option percentage and min_total_reads set to 0), spark.vc (with covariates set to NULL and library sizes set to the total number of counts per sample) and spark.test. SPARK-X was run with function sparkx (option set to “mixture”). MERINGUE was run using functions normalizeCounts, getSpatialNeighbors, and getSpatialPatterns. Seurat’s FindSpatiallyVariableFeatures was run with assay set to “Spatial”. Giotto was applied using functions createGiottoObject, normalizeGiotto, createSpatialNetwork, and binSpect (with bin_method set to “kmeans” or “rank” and got_av_expr and get_high_expr set to FALSE).

### Application to large collection of bulk RNA-seq data

We downloaded a large collection of RNA-seq samples covering 76 mouse cell types and tissues ^23^. This data has been normalized using Upper Quartile normalization and treated for batch effects using ComBat ^35, 36^. From this dataset, we selected samples obtained from brain, prefrontal cortex, hippocampus, cortex, frontal cortex, olfactory bulb, cerebellum, forebrain, neocortex, and cerebral cortex. We treated the data using Seurat, including scaling, finding 1,000 highly variable genes, and PCA. We filtered out genes with generally low expression (mean expression in the bottom 25%), and applied singleCellHaystack on the resulting 1,958 samples and 18,260 genes, using as input space the first 5 PCs, using 25 grid points and otherwise default parameters. For visualization purposes, we applied UMAP on the first 5 PCs.

### Application to module scores of gene sets

We used the R package msigdbr (version 7.5.1) to retrieve sets of mouse genes associated with BIOCARTA pathways ^32^. For 292 pathways which had at least 10 associated genes, we collected their genes, and used the Seurat function AddModuleScore to calculate module scores in the spots of the Visium datasets. These module scores reflect the general activity of the genes in each pathway. Subsequently, we ran singleCellHaystack on each Visium datasets, using as input the coordinates of spots and the module scores of all pathways. High-scoring pathways thus reflect pathways with spatial differences in activity within the tissue (see Fig. 5 for examples).

## NOTES

### Code availability

singleCellHaystack is implemented as an R package and is available from CRAN (https://CRAN.R-project.org/package=singleCellHaystack) and GitHub (https://github.com/alexisvdb/singleCellHaystack), and as a Python package available from PyPI (https://pypi.org/project/singleCellHaystack) and GitHub (https://github.com/ddiez/singleCellHaystack-py).

### Data availability

Single-cell RNA-seq datasets analyzed in this study are available from https://figshare.com/articles/dataset/Robject_files_for_tissues_processed_by_Seurat/5821263 (Tabula Muris), https://figshare.com/articles/MCA_DGE_Data/5435866 (Mouse Cell Atlas), https://atlas.fredhutch.org/nygc/multimodal-pbmc/ (PMBC CITE-seq), https://support.10xgenomics.com/single-cell-multiome-atac-gex/datasets/1.0.0/pbmc_granulocyte_sorted_10k (PBMC scRNA-seq and scATAC-seq).

Visium data was obtained through the SeuratData package. Slide-seqV2 and HDST data from the Broad Institute Single Cell Portal, accession numbers SCP815 and SCP420. MERFISH data from the supplementary data in Xia et al. Bulk RNA-seq data was downloaded from https://figshare.com/articles/dataset/Mouse_data/14178425/1.

## Acknowledgments

The authors would like to thank Y. Harada for secretarial assistance. **Funding:** This work was supported by JSPS KAKENHI Grant Numbers JP20K06609 (A.V.) and JP20K07538 (D.D.), and by an Office of Directors’ Research Grant provided by the Institute for Life and Medical Sciences (Kyoto University). The funders had no role in study design, data collection and analysis, decision to publish, or preparation of the manuscript. **Author contributions:** A.V. conceived of the project and methodology. A.V. and D.D. implemented the methods, ran the analyses and wrote the manuscript. All authors contributed to critical revision of the manuscript. **Competing interests:** All other authors declare no competing interests.

