## Supplementary material for "A universal differential expression prediction tool for single-cell and spatial genomics data"

### Supplementary Figures

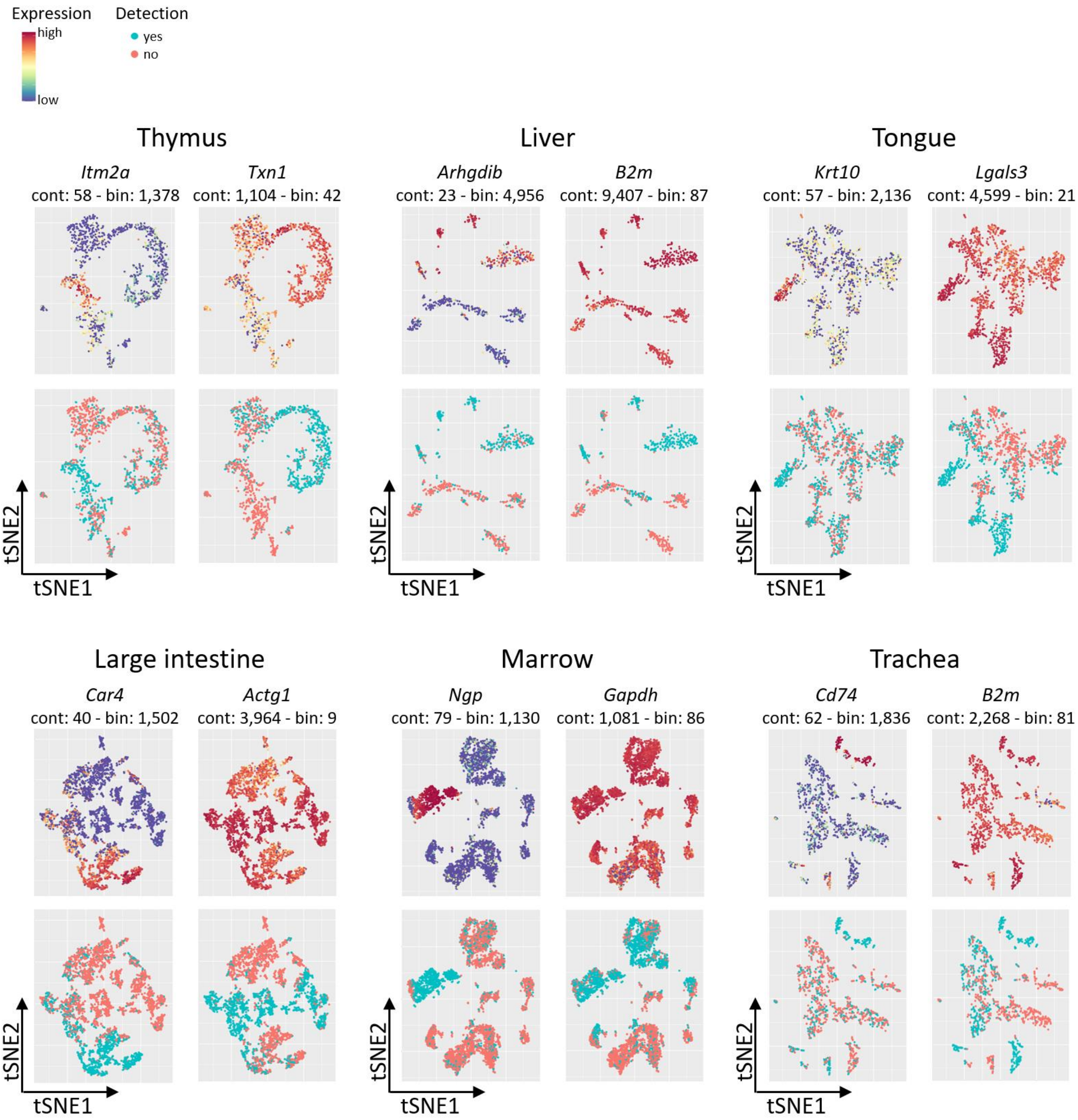


**Supplementary Figure S1:** Additional examples of differences between the original binary method and the new continuous method of singleCellHaystack. For 6 Tabula Muris tissues, examples are shown of genes that are high-scoring according to the new continuous singleCellHaystack but not according to the original binary approach (left side), and vice versa (right side). For each gene, the gene symbol and the ranks according to the continuous (“cont”) and binary (“bin”) approaches are shown, as well as tSNE plots with the continuous expression levels (top) and detection levels as used by the binary approach (bottom).

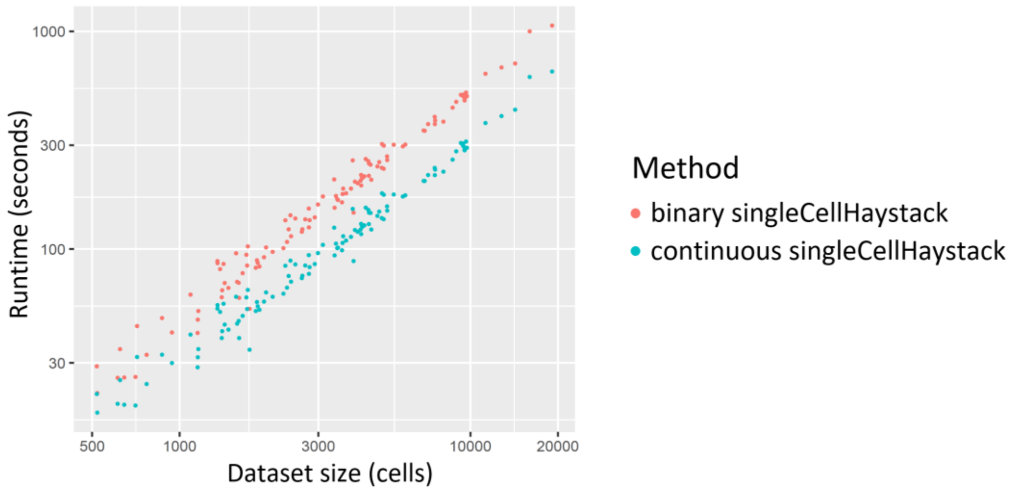


**Supplementary Figure S2:** Comparison of runtimes of the original binary approach (red) and the new continuous approach (blue) of singleCellHaystack on 119 scRNA-seq datasets of Tabula Muris and Mouse Cell Atlas.

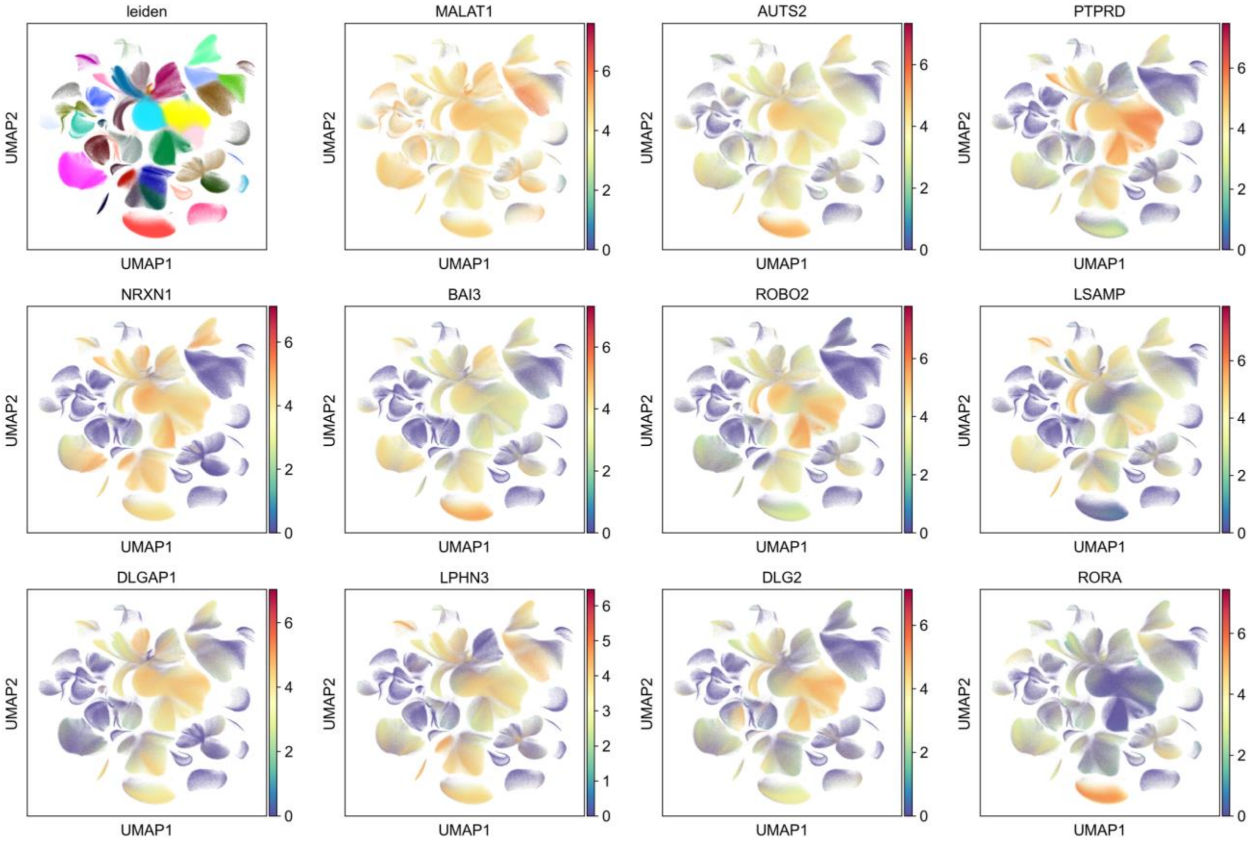


**Supplementary Figure S3:** Application to an atlas-level scRNA-seq dataset. Top 11 genes identified with singleCellHaystack-py using the 4.3 million cells from the Human Organogenesis Cell Atlas using 50 PCA coordinates.

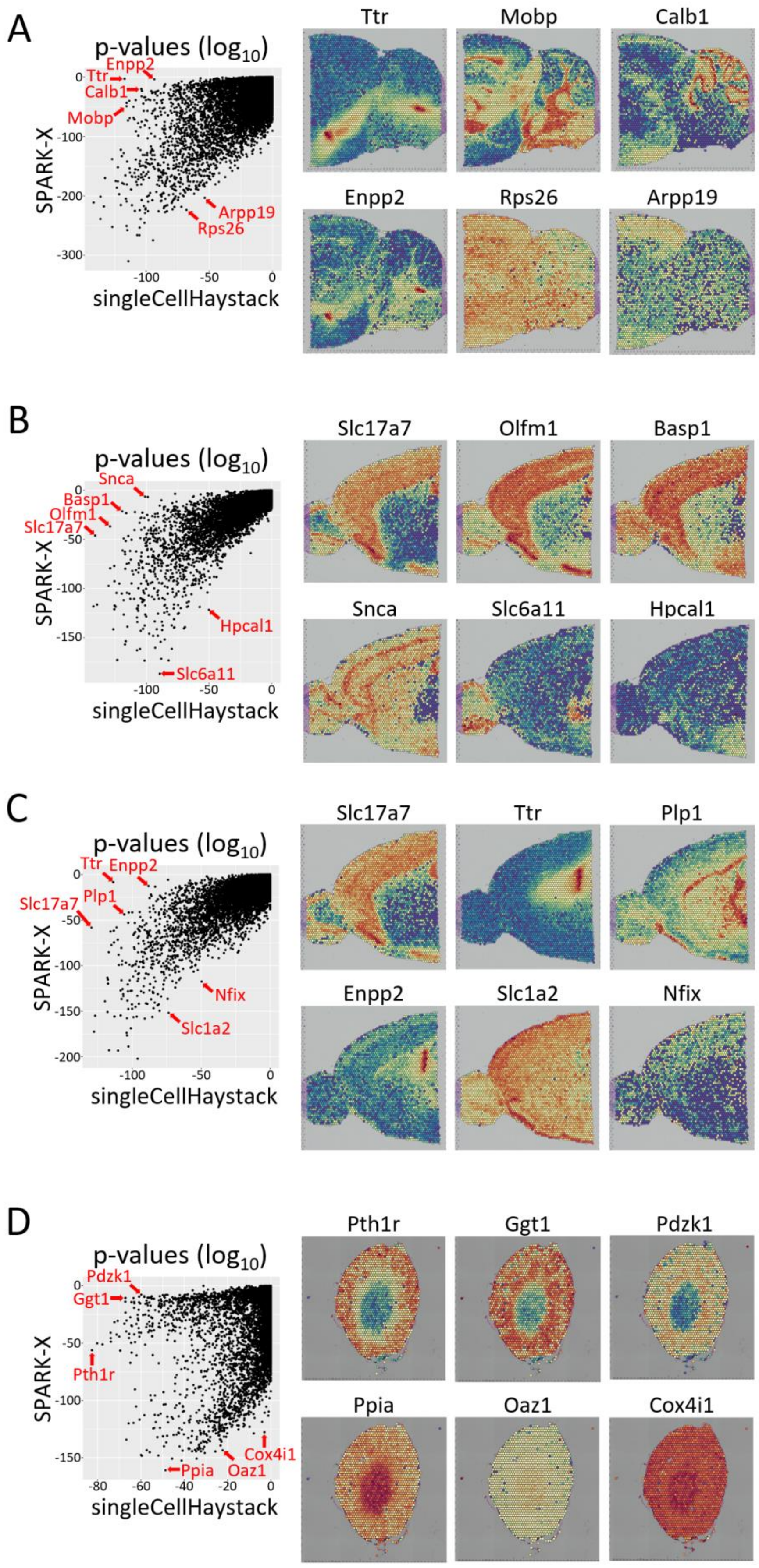
A
p-values ($\log_{10}$)
SPARK-X
singleCellHaystack
Ttr
Enpp2
Calb1
Mobp
Arpp19
Rps26
Ttr
Mobp
Calb1
Enpp2
Rps26
Arpp19
B
p-values ($\log_{10}$)
SPARK-X
singleCellHaystack
Snca
Basp1
Olfm1
Slc17a7
Hpcal1
Slc6a11
Slc17a7
Olfm1
Basp1
Snca
Slc6a11
Hpcal1
C
p-values ($\log_{10}$)
SPARK-X
singleCellHaystack
Ttr
Enpp2
Plp1
Slc17a7
Nfix
Slc1a2
Slc17a7
Ttr
Plp1
Enpp2
Slc1a2
Nfix
D
p-values ($\log_{10}$)
SPARK-X
singleCellHaystack
Pdzk1
Ggt1
Pth1r
Ppia
Oaz1
Cox4i1
Pth1r
Ggt1
Pdzk1
Ppia
Oaz1
Cox4i1

*(previous page)* **Supplementary Figure S4:** Comparison of results of singleCellHaystack and SPARK-X on four 10x Visium datasets. This figure supplements Figure 2B in the main paper. For each comparison, a scatterplot of p values ($\log_{10}$) is shown on the left, and examples of DEGs are shown on the right. For each dataset, DEGs that are high-scoring according to one method but not the other are picked up. Datasets are posterior brain (“posterior2”) **(A)**, anterior brain (“anterior1” and “anterior2”) **(B-C)**, and kidney **(D)**.

*(next page)* **Supplementary Figure S5:** Comparison of results of singleCellHaystack and SPARK-X on four Slide-seqV2 datasets. This figure supplements Figure 2C in the main paper. For each comparison, a scatterplot of p values ($\log_{10}$) is shown on the left, and examples of DEGs are shown on the right. For each dataset, DEGs that are high-scoring according to one method but not the other are picked up. Datasets are from hippocampus **(A)**, embryo **(B)**, olfactory **(C)**, and cortex **(D)**.

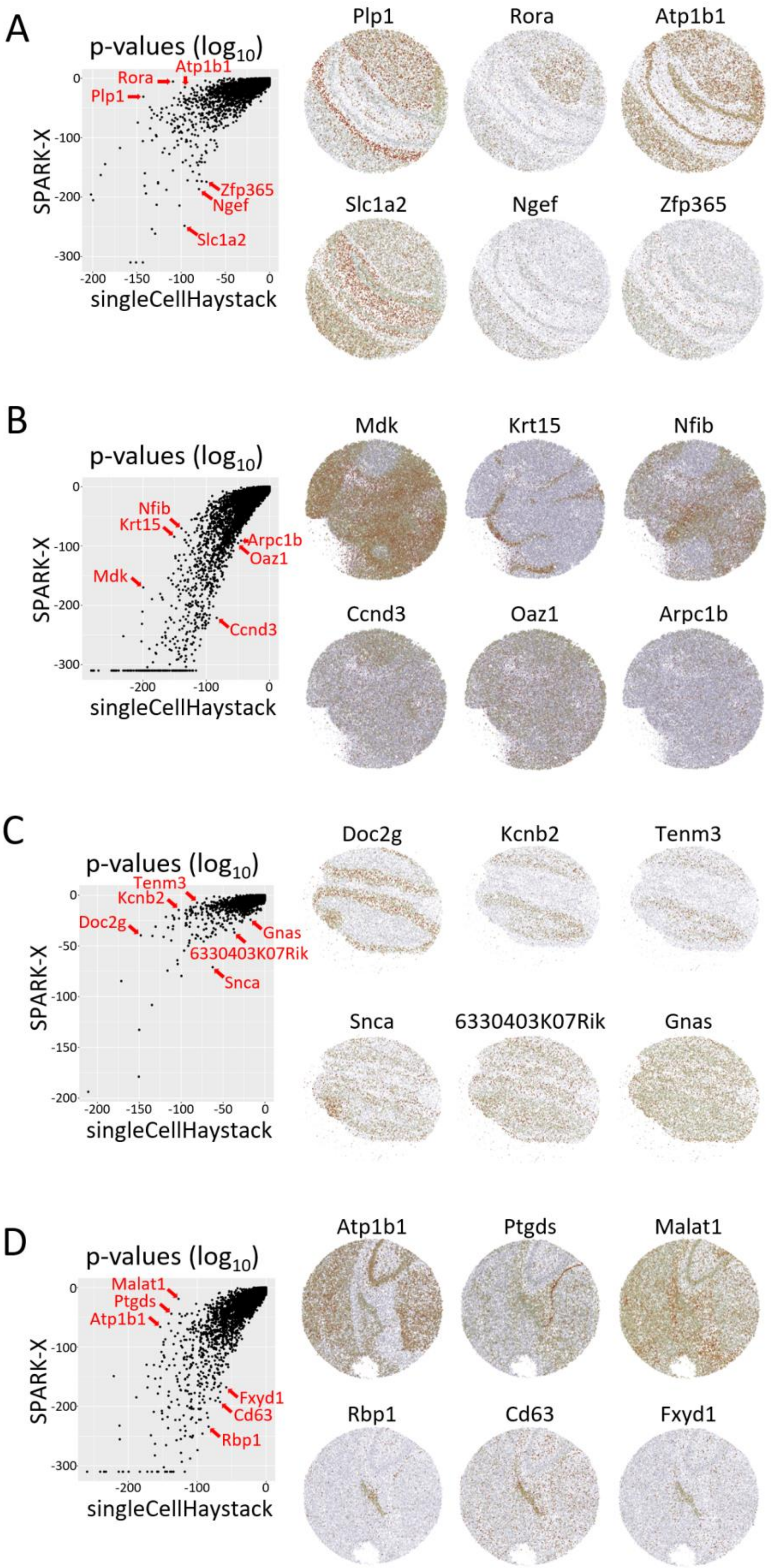

A
p-values (log10)
Rora
Atp1b1
Plp1
Zfp365
Ngef
Slc1a2
SPARK-X
singleCellHaystack
0
-100
-200
-300
-200
-150
-100
-50
0
Plp1
Rora
Atp1b1
Slc1a2
Ngef
Zfp365
B
p-values (log10)
Nfib
Krt15
Arpc1b
Oaz1
Mdk
Ccnd3
SPARK-X
singleCellHaystack
Mdk
Krt15
Nfib
Ccnd3
Oaz1
Arpc1b
C
p-values (log10)
Tenm3
Kcnb2
Doc2g
Gnas
6330403K07Rik
Snca
SPARK-X
singleCellHaystack
Doc2g
Kcnb2
Tenm3
Snca
6330403K07Rik
Gnas
D
p-values (log10)
Malat1
Ptgds
Atp1b1
Fxyd1
Cd63
Rbp1
SPARK-X
singleCellHaystack
Atp1b1
Ptgds
Malat1
Rbp1
Cd63
Fxyd1

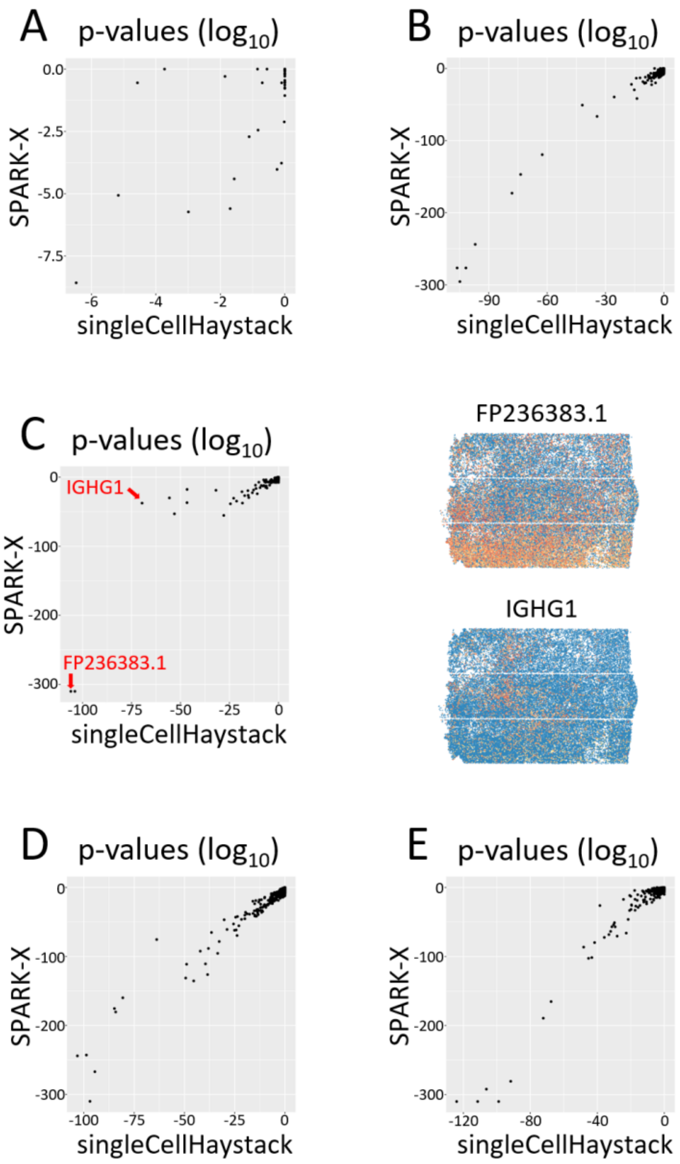


**Supplementary Figure S6:** Comparison of results of singleCellHaystack and SPARK-X on five HDST datasets. This figure supplements Figure 2D in the main paper. For each dataset, a scatterplot of p values ($log_{10}$) is shown. In general, results returned by both methods are very consistent. In dataset vickovic_CN13_D2 **(A)** neither method found any DEGs. In datasets vickovic_CN21_C1 **(B)**, vickovic_CN21_D1 **(C)**, vickovic_CN21_E2 **(D)**, and vickovic_CN24_E1 **(E)**, high-scoring genes of one method were also high-scoring genes according to the other method. For vickovic_CN21_D1 **(C)**, two genes are indicated as an illustration. Both are high-scoring according to both methods.

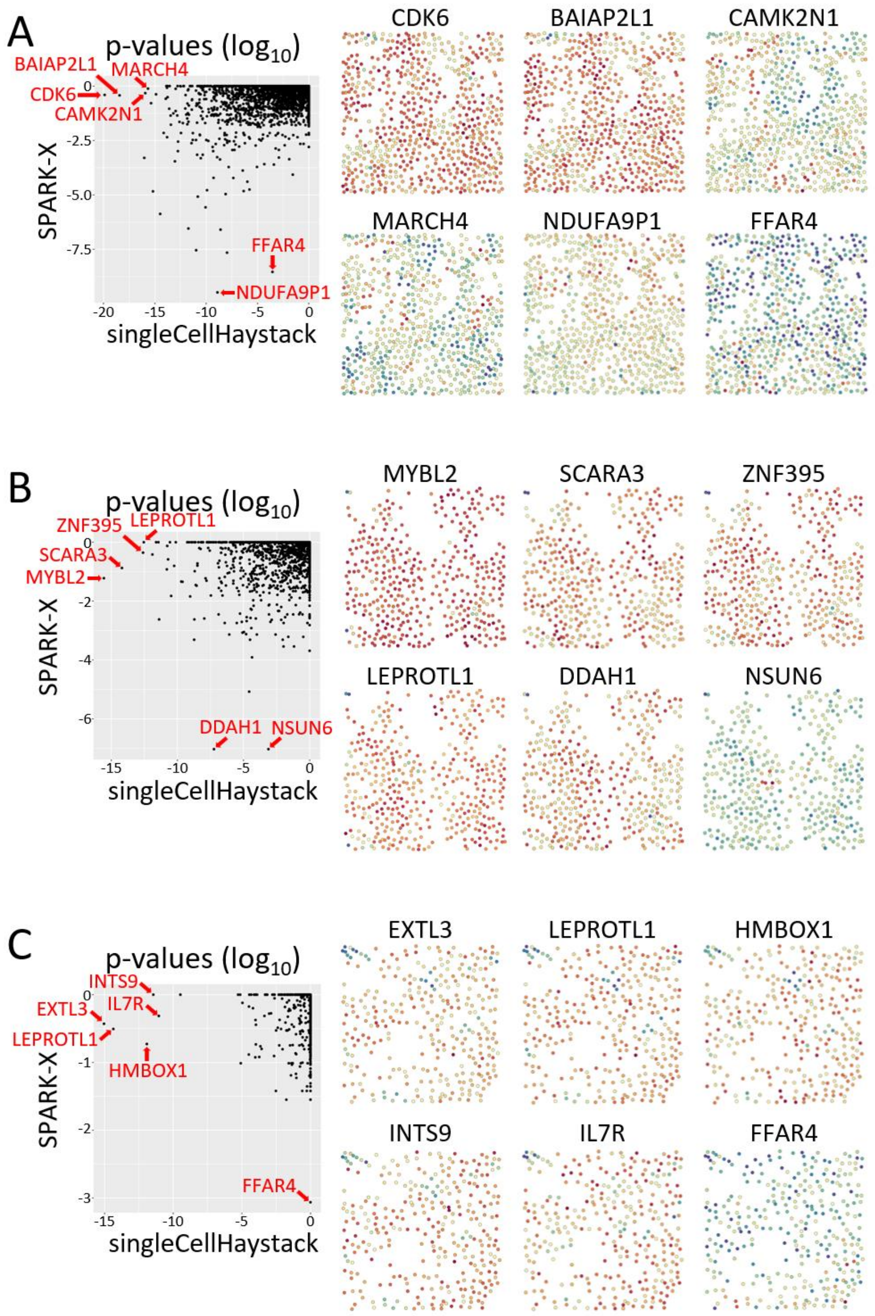


**Supplementary Figure S7:** Comparison of results of singleCellHaystack and SPARK-X on three MERFISH datasets. For each comparison, a scatterplot of p values ($\log_{10}$) is shown on the left, and examples of DEGs are shown on the right. For each dataset, DEGs that are high-scoring according to one method but not the other are picked up. Datasets are Xia *et al.* B1 **(A)**, B2 **(B)** and B3 **(C)**.

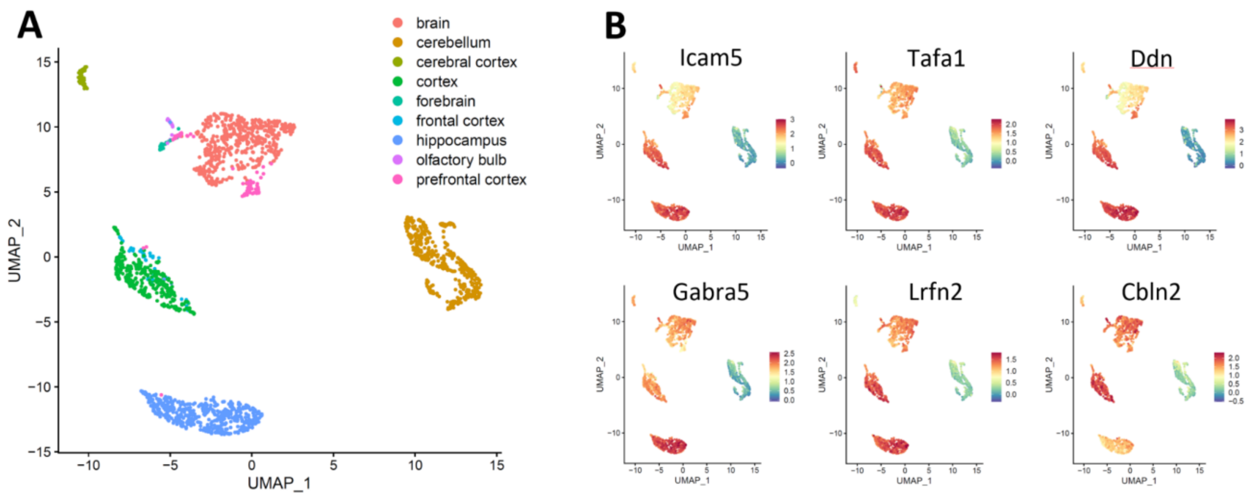


**Supplementary Figure S8:** Application of singleCellHaystack on a collection of bulk RNA-seq samples. **(A)** UMAP plot of the 1,958 RNA-seq samples obtained from various parts of the mouse brain. **(B)** The top 6 high-scoring DEGs predicted using singleCellHaystack.
